# Fast Cycling Culture of the Marine Annelid *Platynereis dumerilii*

**DOI:** 10.1101/2023.04.30.538804

**Authors:** Mathieu Legras, Giulia Ghisleni, Rabouant Soilihi, Enzo Celmar, Guillaume Balavoine

**Affiliations:** Université de Paris Cité, CNRS, Institut Jacques Monod, F-75006 Paris, France; University of Milano-Bicocca, Department of Biotechnology and Biosciences

## Abstract

*Platynereis dumerilii*, a marine annelid, is a model animal that has gained popularity in various fields such as developmental biology, biological rhythms, nervous system organization and physiology, behaviour, reproductive biology, and epigenetic regulation. The transparency of *P. dumerilii* tissues at all developmental stages makes it easy to perform live microscopic imaging of all cell types. In addition, the slow-evolving genome of *P. dumerilii* and its phylogenetic position as a representative of the vast branch of Lophotrochozoans add to its evolutionary significance. Although *P. dumerilii* is amenable to transgenesis and CRISPR-Cas9 knockouts, its relatively long and indefinite life cycle, as well as its semelparous reproduction have been hindrances to its adoption as a reverse genetics model. To overcome this limitation, an adapted culturing method has been developed allowing much faster life cycling, with median reproductive age at 15 weeks instead of 6-8 months using the traditional protocol. A low worm density in boxes and a strictly controlled feeding regime are important factors for the rapid growth and health of the worms. Moreover, a genetic selection for fast-reproducing individuals has been applied to isolate a “Fast Forward” strain that can be used for egg microinjection. This culture method has several advantages, such as being much more compact, not requiring air bubbling or an artificial moonlight regime for synchronized sexual maturation, and necessitating only limited water change. A full protocol for worm care and handling is provided.

## Introduction

During the process of experimental design, the choice of the most appropriate model organism is a crucial step that determines the validity of the study. Gregor Mendel, the pioneering author of some of the most famous genetics experiments on plants of the genus *Pisum*, stated that the selection of an unsuitable model can make the results questionable from the outset (Abbott & Fairbanks, 2016). The pea plant was not only physiologically and morphologically suitable for his “Experiments on Plant Hybrids”, but its representational power has made it a model organism by definition, (Ankeny & Leonelli, 2020; Irion & Nusslein-Volhard, 2022). Knowledge derived from a model organism can be projected and generalized to a broader range of systems, and this, together with the integration of multidisciplinary approaches, allows for a 360-degree comparative study between species across the tree of life. For example, in the case of bilaterian animals, studying mice, chickens, fish, and sea urchins (which belong to the deuterostomes) together with *C. elegans* and *D. melanogaster* (from the ecdysozoans group) has enabled a comparative approach to regulatory cladistics that has revealed the shared nature of signalling pathways and transcription factors among all modern Bilaterians (Pires-daSilva & Sommer, 2003). The evolution of phylum-specific body plans and morphological structures is generally not due to the rise of new developmental genes, but we can expect and predict that novel regulation circuits have developed (Peterson & Davidson, 2000). Investigating how developmental genes are regulated in different lineages can therefore provide new evolutionary insights into the beginnings of the history of Bilaterians.

In this context, the marine annelid *Platynereis dumerilii* (Özpolat et al., 2021) compensates for the longstanding lack of representatives of the lophotrochozoans, which is the third branch of bilaterians, alongside deuterostomes and ecdysozoans, that has impeded a reliable approach to bilaterian comparative genetics. Carl Hauenschild was the first scientist to begin culturing *P. dumerilii* in 1953 and standardize its culture conditions (Hauenschild & Fischer, 1969). The potential of *P. dumerilii* as an emerging animal model lies in the study of its molecular and developmental processes. It is considered to have retained some ancestral characteristics and, as a polychaete annelid, to have only slightly deviated from *Urbilateria*, the last common ancestor of all Bilaterians (Balavoine, 2014; Tessmar-Raible & Arendt, 2003). Earthworms and leeches have been used as lophotrochozoan animal models, but they show derived features within the annelid clade. They lack many of the head structures and trunk appendages, and their life cycle lacks a larval phase and metamorphosis, which are ancestral developmental traits of annelids (Struck et al., 2014). In contrast, all these characteristics are present in the polychaete family nereididae, to which *P. dumerilii* belongs, making it a suitable and strategic model for providing insight into the deep morphological and developmental past of all bilaterians.

*P. dumerilii* is found in all European seas. When managing hundreds or thousands of individuals, the fact that they are undemanding, small-sized, and easily manipulated is crucial. Regarding its life cycle, a zygote can become a sexually mature adult in a few months under culture conditions, or about 1 year in the wild. The ability to culture *P. dumerilii* in the laboratory for the full life cycle, its rapid and highly synchronized external embryogenesis, and the transparency of embryos and juveniles (which allows for live imaging and the use of optogenetic tools) represent a significant advantage for studying reproductive biology, development, and regeneration.

The life cycle stages, and the development have already been well described (A. Fischer & Dorresteijn, 2004; A. H. L. Fischer et al., 2010; Hauenschild & Fischer, 1969; Hempelmann, 1911). The following description is just a summary, with stages given at a culturing temperature of 18°C. A single mating of *P. dumerilii* can result in thousands of eggs (160 µm diameter). Once fertilized, the eggs develop into larval stages in a synchronized manner. The zygote contains lipid droplets, cortical granules, protein yolk granules, and secrete a protective jelly to avoid polyspermy (A. Fischer & Dorresteijn, 2004) and protect the embryo from predation. Copious jelly secretion indicates successful fertilization. During the early stages of development, *P. dumerilii* embryos exhibit spiral cleavage. The first divisions are highly asymmetric, and cells can be identified based on their position and size. During this spiral phase, the lipid droplets fuse to form four large lipid droplets, which indicate healthy development. Larvae with more or less than four lipid droplets will not develop normally.

By 24 hours post-fertilization (hpf), the trochophore larvae are free from their jelly, start swimming with the help of a ciliary belt, and exhibit positive phototaxis. At this stage, larvae are planktonic and do not feed. The only food source is the yolk contained in their macromeres. By 4 days, the nectochaete larvae have produced a head, a pygidium (posterior end), tentacular cirri, anal cirri, and three chaetigerous segments with parapodia. The larvae then move from the pelagic zone to the benthic zone and create a mucus tube to spend most of their life in it. At 6 days, they begin feeding (mostly at night when they leave the tube) and add new segments to the posterior end from the segment addition zone (SAZ), located just anterior to the pygidium (Balavoine, 2014; A. H. L. Fischer et al., 2010). From now on, the developmental pace of juveniles will vary between individuals. The size of the worm depends on its food regime and quantity of food intake. Before sexual maturation, gonial clusters begin to populate the body of the worms when they reach around 40 segments (Kuehn et al., 2022). Gametes start to populate the coelom at approximately 50 segments in length but become mature at ∼70-80 segments in size (A. Fischer, 1974, 1975). During sexual metamorphosis, the worms stop feeding and their gut degenerates. Their eyes increase in size and extensive changes occur in their muscles, parapodia, and body color, as they transition from a sexually immature atoke form to a sexually mature epitoke form. Sexually mature worms need to swim very fast, so their muscles degenerate to make room for a new epitokous muscle type. Oocytes are yellow and give this color to mature females. Males are bicolor due to the white sperm that colors the anterior part of the worm, while the posterior part is red due to blood capillaries filled with hemoglobin (Andreatta et al., 2020; A. H. L. Fischer et al., 2010). Once epitoky is complete, spawning of *P. dumerilii* peaks around 1 week after the full moon phase (Zantke et al., 2013). The worms leave their tubes and move into pelagic water to look for a partner. Aroused by pheromones (Zeeck et al., 1991, 1998), the mating couple starts a nuptial dance (circular fast swimming) and the female releases all of her oocytes into the water. The male fertilizes them externally and the worms die shortly after the release of gametes. All of these developmental stages can be influenced by changes in temperature (A. Fischer & Dorresteijn, 2004).

Several studies have demonstrated the suitability of *P. dumerilii* for reverse genetics experiments, including transgenesis using DNA transposase (Backfisch et al., 2013, 2014) and targeted gene knockouts with TALENs or Cas9 (Bannister et al., 2014; Bezares-Calderón et al., 2018). However, despite its potential as a genetic model organism, there are still challenges to making *P. dumerilii* a widely used “fruit fly of the sea.” One of these challenges is the extensive genetic polymorphism present in laboratory strains, even after decades of maintenance in various European and American laboratories. Non-inbred strains still exhibit a high proportion of SNPs (Single Nucleotide Polymorphism). This is particularly relevant to genome editing by CRISPR-Cas9, as a PCR polymorphism study must be conducted for each targeted gene. Furthermore, reproduction in P. dumerilii occurs only once at the end of the life cycle, and adults die immediately after spawning. This makes it difficult to maintain homozygous strains, as adults of both sexes must be obtained on the same day, requiring large batches of transgenic animals to be produced and maintained until they reach sexual maturity.

The main challenge remains the relatively long reproductive cycle of *P. dumerilii*. In nature, the reproduction period is seasonal, implying that most worms live for about a year on average. In the laboratory, temperature (generally 18°C) and light regimes (8/16 hrs of night/day) are used to simulate the end of springtime. To replicate the role of moonlight in synchronizing the swarming of male and female adults, an artificial moon, usually in the form of a small bulb, is placed in the culture room for one week every four weeks. These conditions, combined with regular feeding, sometimes induce sexual maturation at a much younger age. However, the age of reproduction remains highly variable and is not correlated between worms from the same parental batch. A typical culture containing around 30 juvenile worms will usually produce adults whose ages vary between 4 and 12 months, making it difficult to create and maintain transgenic strains.

We hypothesize that three factors contribute to the variability in sexual maturation age. Firstly, the highly variable density of worms in culture boxes may be a factor. *P. dumerilii* juveniles are typically kept in flat plastic alimentary boxes with a bottom surface area of around 500 cm^2^. Young worms spin silk tubes at the bottom of the box in which they spend most of the day. Instead of being randomly distributed, the tubes are usually built in positions that maximize distances between worms, suggesting territorial behaviour. This behaviour is likely induced by antagonistic interactions between juvenile worms, which can be observed frequently as wandering worms are attacked and bitten by their neighbours. Secondly, the quantity of food delivered is not adapted to the number of worms in the boxes. Too little food results in slow growth, while too much food can lead to water fouling with the same effect or even worm death by asphyxia. In classical culture methods, water must be changed every two weeks to tackle fouling. Finally, the extensive genetic polymorphism observed in our historical culture population (which we call the polymorphic strain) and verified in numerous PCR studies on various genes has led us to hypothesize that genetic factors may also be involved in the variability of maturation age.

In this article, we describe how we have established a new protocol for culturing a selected strain of *P. dumerilii* called Fast Forward (FF) by controlling density, adapting food delivery, and selecting worms based on their maturation age. Sexual maturation for this strain begins at 12 weeks, and most worms mature before 18 weeks. This protocol places *P. dumerilii* among the small minority of animal models in which transgenic and genome editing techniques can be developed with ease in the future. Most importantly, we have significantly simplified the culture conditions by eliminating the need for a moon cycle and reducing the frequency of water changes, thereby reducing staffing requirements and enabling the rearing of several transgenic strains in limited lab space.

## Materials and Methods

### Making new batches of larvae

Swimming adults are collected from low density boxes every morning. Male worms display a bi-coloured white/red pattern, whereas female worms are yellow/orange. Only adults actively swimming are chosen daily for reproduction. To make all fertilizations and keep the embryos/larvae up to ten days after fertilization, small beakers (diameter 9.5 cm, height 5.5 cm) are used. Approximately 150 ml of NFSW is added in each beaker, along with one pair of worms. All worms are manipulated using plastic pipettes (Samco scientific). Ideally, the release of gametes occurs in the next few minutes. Sometimes, this release does not happen spontaneously even though both individuals are active. In such cases, spawning can be achieved by gently pressing the male with a fine paintbrush to force the release of a small amount of sperm. This, in turn, triggers the release of the female oocytes. Once both adults have released their gametes, most of the NFSW used for fertilization must be poured out to prevent overnight fouling by the male sperm. This is done easily since the fertilized eggs sediment quickly at the bottom of the beaker. A new volume of 150 ml of clean NFSW is then added. The efficiency of the fertilization is checked by the formation of a transparent jelly coat around the eggs. Thirty minutes post fertilization, the eggs adopt a characteristic hexagonal spatial distribution while being pushed apart by the jelly. To prevent fouling of the water due to the decomposition of the jelly coat, 1.5 ml of 100x penicillin:streptomycin mix is added. Fertilized batches are incubated overnight at 20°C. The next day, at 24 hours post-fertilization (24 hpf), the larvae must be dejellified (getting rid of the jelly coat). The content of the beaker is poured into a large 80 μm sieve (see lab-made equipment section), allowing the jelly to sieve through with the seawater. After two quick rinses with NFSW, the sieve is returned upside down to the original beaker and 150 ml NFSW is poured over the whole sieve to recover all swimming larvae. Small worms are incubated at 20°C until they reach 4 dpf. They are then fed with 3 ml of 1x frozen microalgae (*Tetraselmis marina*, Instant Algae™, Table 1). Three-segment small worms should start to feed and settle at the bottom of the beaker quickly.

**Table 1.** Weekly schedule for the distribution of food in individual boxes. All volumes are suspensions in NFSW at the concentration indicated in the text. Worm ages are in days post fertilization (dpf).

| Worm ages | monday | wednesday | friday |
| --- | --- | --- | --- |
| 4-30 dpf | 3 ml algae |  | 3ml algae |
| 31-45 dpf | 1ml Sera micron® | 3 ml spinach | 1ml Sera micron® |
| > 46 dpf | 3ml Tetramin® | 3ml spinach | 3ml Tetramin® |

### Transplanting new boxes of juvenile worms

To achieve maximum survival of juvenile worms, it is important to use healthy batches of young worms. At 10 dpf, the beaker containing young worms should consist primarily of feeding individuals, which can be checked by a gut full of algae and the budding of a fourth segment in some. For further culturing, we will use these healthy and rapidly growing 4-segment young worms and avoid 3-segment worms and those with an empty gut. The boxes used for the main phase of growth are polypropylene containers, that are 6.5 cm high, and can be easily stacked in incubators or on the bench. These containers have lids that are not completely airtight, ensuring vital gas exchange for the worms. Typically, four boxes are transplanted from each batch of young worms. Each box is filled with 500 ml of NFSW and supplemented with 3ml of 1x frozen algae and one cm^2^ of old box algal mat (see commensals section). Twenty young worms are transplanted into each box, achieving the optimal density of 300 individuals.m^-2^. Young worms can be selected using a P20 hand pipette with bevelled pipette tips. Worms reflexively grip the inside of the tip with their bristled appendages, so it is important to set the hand pipette at a very small volume (1 μl) to pipette individual worms and expel them out of the tip efficiently. The boxes are incubated at 20°C, with a 16-hour day / 8-hour night light regime, for the entire remaining life cycle.

### Food regime

The food regime is inspired from the classical culture system (Hauenschild & Fischer, 1969). Four different types of food are used successively to ensure rapid and healthy growth. The quantities mentioned must be strictly followed, as overfeeding can cause water fouling and be a major cause of death for juvenile worms. The microalgae typically used for feeding early juvenile stages are live *Tetraselmis marina*. To avoid time consuming lab culture of this green alga, commercial fish hatchery food (Instant Algae™, Reed Mariculture) is used. The algae are diluted 40 times in NFSW, aliquoted in 50 ml Falcon tubes and kept frozen at -20°C. Sera Micron® (Sera) is the food chosen for older worms, as its particle diameter is well suited for the worm’s mouth at this stage and it is highly nutritious for rapid growth. It is resuspended (1% w/v) in NFSW and must be kept frozen if not used immediately. Tetramin® flakes are fed to sub-adults. They are ground to a powder using a mortar and are suspended in NFSW (1% w/v). Tetramin® suspension can be kept frozen if not used on the same day. Finally, frozen organic spinach is used to supplement both Sera® and Tetramin® regimes, in a 10% w/v suspension that is ground with a food blender. Food distribution is made according to table 1.

### Worm growth monitoring and water change

Young worms are not visible to the naked eye during the first three weeks of culturing in polypropylene boxes. After 3-4 weeks, they start spinning silk tubes that are large enough to be observed at the bottom of the containers. Therefore, it is recommended to check after one month that growth is normal, and no mortality of young worms has occurred. At two months (61 days), a water change is necessary. At this age, worms are firmly settled in their silk tubes during the day and the old water can be safely poured out into an intermediate container before being discarded. Occasional runaway worms can be put back into their box with a plastic Pasteur Pipette. 500 ml of fresh NFSW is added to each box. Worms are counted at this 2-month stage to ensure that there are 20 worms or near to 20 worms in each box. As the worms will sexually mature rapidly over a period of 4-5 weeks starting at 12 weeks, no further water change will be required. As the worm counts decrease in each box through maturation, boxes containing five remaining worms or fewer are regrouped to save food and space in the incubator. To handle worms out of their box, plastic Pasteur pipettes are used. Worms are chased from their tube by gently compressing the silk tube with the pipette tip, starting at the end of the tube where the head of the worm is facing. In this way, the worm wiggles backward out of the tube, reducing the risk of occasional injury and sectioning of the fragile tail.

### Commensals

*P. dumerilii* cultures are anything but axenic. Boxes contain several other organisms, some of which are likely beneficial for the growth and maintenance of the annelid, while others may grow to the point of interfering with the well-being of the worms. These co-cultured organisms are typically passed down from generation to generation when transplanting worms to new boxes with pipettes and are not detected due to their microscopic size. Alternatively, they also come with the natural sea water (in our case, filtered natural sea water from the bay of Cancale in Brittany), which, despite filtering, is not sterile. It is important to mention that we have never observed cases of internal parasitism, such as by myxozoans (Rangel et al., 2009), in our culture. This indicates that the filtration quality of the sea water we used has been satisfactory in this respect. In this paragraph, we will give a brief description of these commensals. We will describe how to transfer them from boxes to boxes if they may be a complementary source of food for the worms, help clean out excess distributed food, or help maintain the homeostasis of the box in any other respect. We will also describe how to get rid of unwanted commensals.

In the older boxes, the bottom gradually becomes covered with green filamentous algae and cyanobacteria mats (figure 3). Small worms feed on filamentous algae, as the algae gets cleared near the extremities of the worm tubes. Additionally, these photosynthetic organisms are likely important in maintaining a satisfactory level of dissolved oxygen in the water, as we never place air diffusers at any stage of this culture. We have observed that small worms settle in tubes earlier in boxes where we ensure that filamentous algae and other components of the mat grow rapidly. To obtain quick growth of the mat, we inoculate boxes with small pieces of the mat from a 4-month-old box. It is important to ensure that this old box does not contain any unwanted commensals (such as *Dimorphilus* worms, see below) or small *P. dumerilii* larvae or tiny juveniles from spontaneous reproduction, which can occur even when boxes are checked for adults daily. The mat is scraped from the box bottom using a plastic scraper. A small piece roughly 1 cm^2^ is sufficient to inoculate a new box. This is done at the same time as new boxes are populated with little 10-day worms (figure 2).

**Figure 1.**
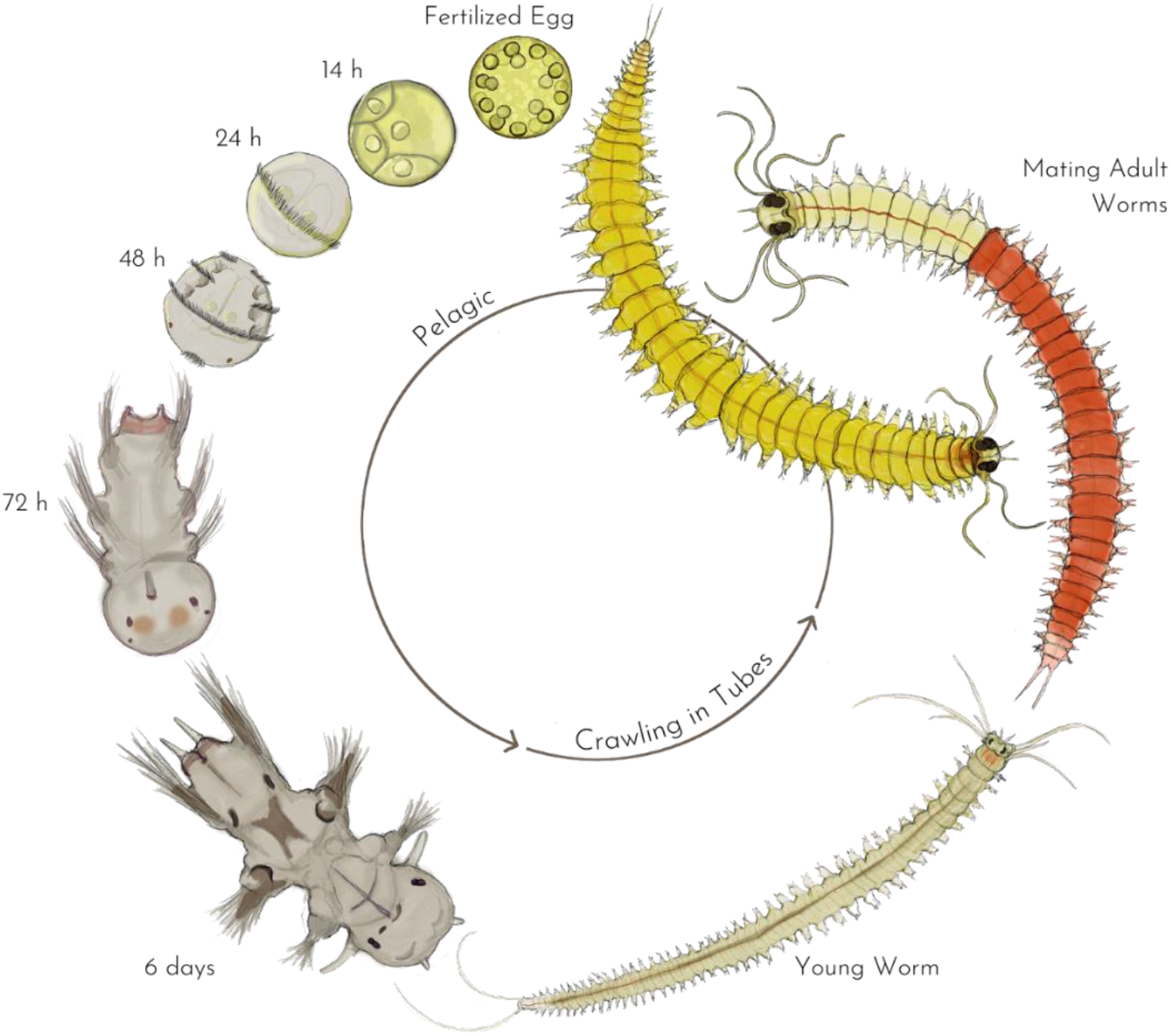
*Platynereis dumerilii*’s life cycle. The different life stages are represented and categorized according to the pelagic/benthic behavior of the worm. From the fertilized egg to the 48 hpf stage, the organism has a diameter of about 160 µm. In the 72h stage, the swimming larva is about 250 µm long, while at 6 days it reaches about 400 µm (A. H. L. Fischer et al., 2010).

**Figure 2.**
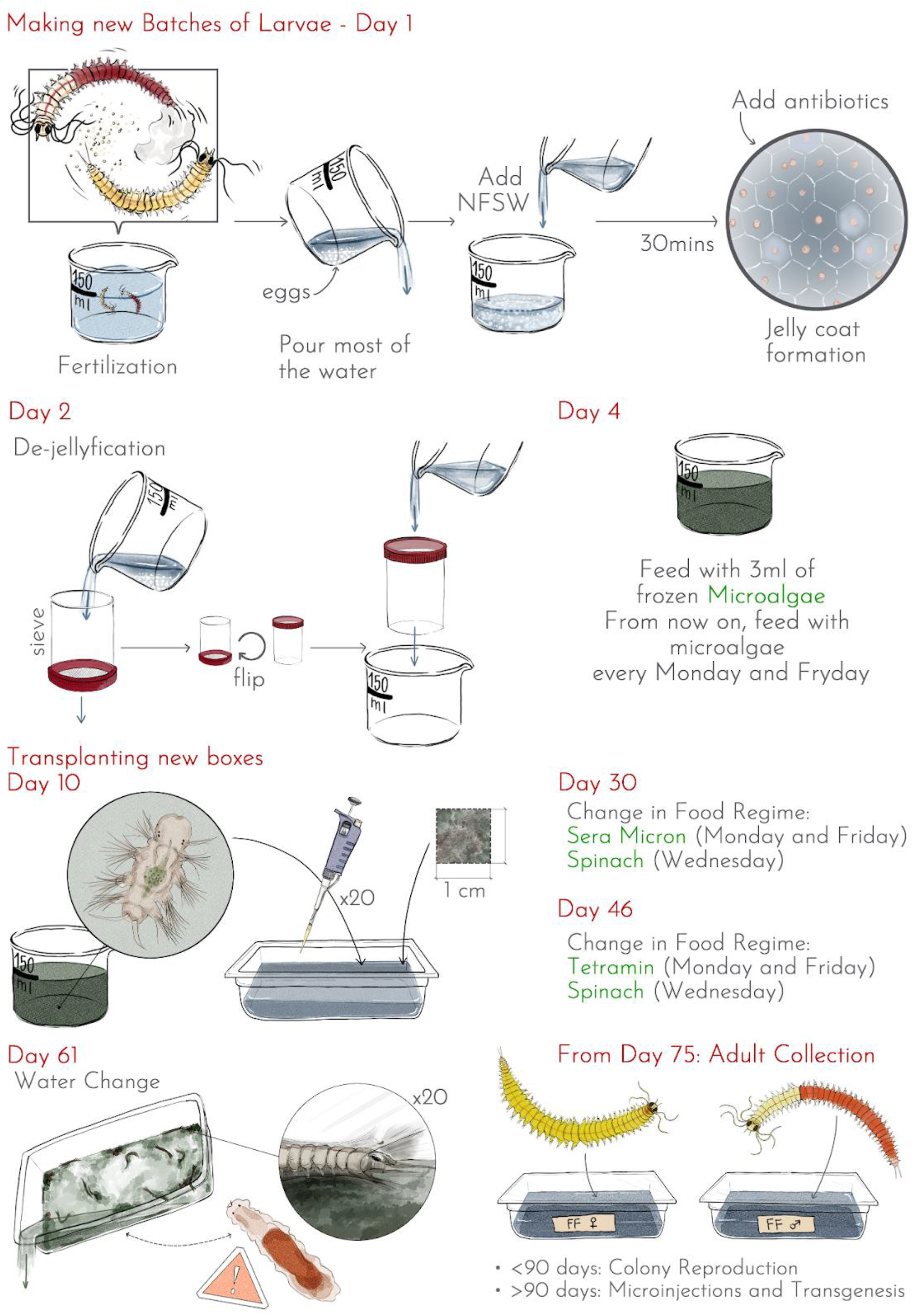
*Platynereis dumerilii* Fast Forward strain Protocol overview. The overall progression to make the selection of FF individuals and reach the proposed protocol is described in the Results section.

**Figure 3.**
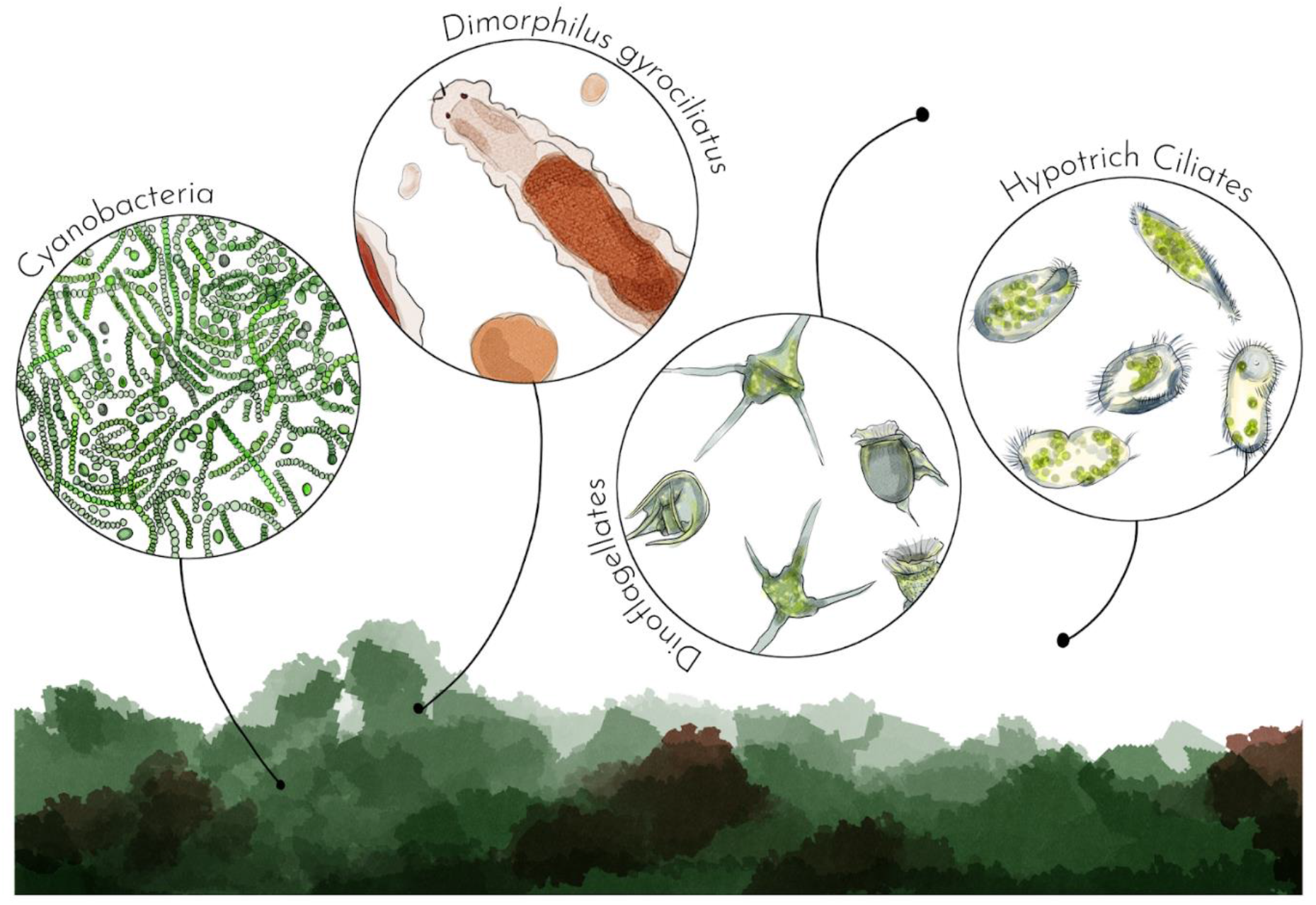
Commensals commonly found in *Platynereis dumerilii*’s culture boxes. Cyanobacteria and filamentous green algae are constitutive of the mat. *Dimorphilus*, nematodes and ciliates feed at the surface of the mat. Dinoflagellates are free swimming above the mat.

Unicellular eukaryotes are also present in our culture boxes, consisting of two types: hypotrich ciliates wandering on the box bottom and swimming flagellated unicells (mostly dinoflagellates). In boxes, their proliferation is harmless to the worms and may even benefit the culture by cleaning out excess food. These unicellular eukaryotes are propagated from one generation of boxes to the next by pipetting adults or seeding filamentous algae, as practiced in the lab. These organisms do not require extensive monitoring, except for one exception. In the beakers containing larvae and small worms fed with *Tetraselmis*, dinoflagellates can proliferate excessively, rendering the water cloudy, depleting the medium of oxygen, and ultimately killing the small worms. If the recommended quantity of food and culture time in these small beakers with a high density of worms is followed (no more than 10 days), this problem is unlikely to become uncontrollable.

Lastly, small metazoans are sometimes co-cultured with *P. dumerilii*. Nematodes have been observed on several occasions, but they never reach high densities and do not affect the wellbeing of *P. dumerilii*. However, more concerning are the proliferations of the tiny annelid *Dimorphilus gyrociliatus* (Martín-Durán et al., 2020). These limbless, gliding annelids, which are less than 1 mm long, can be detected by examining the box bottoms with a stereomicroscope. In some of our low-density culture boxes, they have reached enormous densities, hindering the growth of *P. dumerilii* and potentially causing their death. To eliminate them, we transferred all *P. dumerilii* individuals to a new box. Each worm had to be rinsed with seawater several times to ensure no *D. gyrociliatus* remained. It is also important to follow simple rules to eliminate these unwanted commensals from the culture: the beakers containing the small 10-day worms used to populate new boxes should be checked for any contaminating *D. gyrociliatus*, and plastic pipettes used to collect adults should be discarded and replaced when collecting worms from boxes of a new generation.

### Data analyses and graphs

All analyses were performed in ‘R 4.1.3’ and RStudio. Graphs were created using the R package ‘ggplot2 v3.4.1’.

## Results

### Selection of fast reproducing individuals

An initial selection of four pairs of early-reproducing worms was made from the polymorphic stock. The worms obtained from these four pairs were called FF1G (1st generation). All crosses were made between worms of the same generation, which were further referred to as FF2G, FF3G, and so on. The dates of emergence of all mature worms were systematically recorded in each generation up to 130 days after fertilization. Only the earliest pairs of mature worms of each generation were retained for spawning the next generation. Importantly, we systematically avoided brother/sister crosses. The rationale for this restriction is to preserve as much of the initial polymorphism as possible to prevent the co-selection of deleterious mutations and the rapid selection of sub-optimal combinations of genes in terms of maturation age. Another crucial factor in this process was the selection of fast-growing juvenile worms from 10 dpf batches to populate low-density boxes (see Materials and Methods). From FF5G onward, we exclusively selected the 10-day post-fertilization worms that had developed a fourth segment after six days of feeding with microalgae (figure 2). This is a second level of selection that may have influenced the final gene frequencies of the FF population. Eight generations were obtained in this way over a two-year period. Excluding the problem that affected FF4G, likely due to the food regime (see next section), the median age of maturing worms decreased from generation to generation (Figure 4) but did not improve after FF6G.

**Figure 4.**
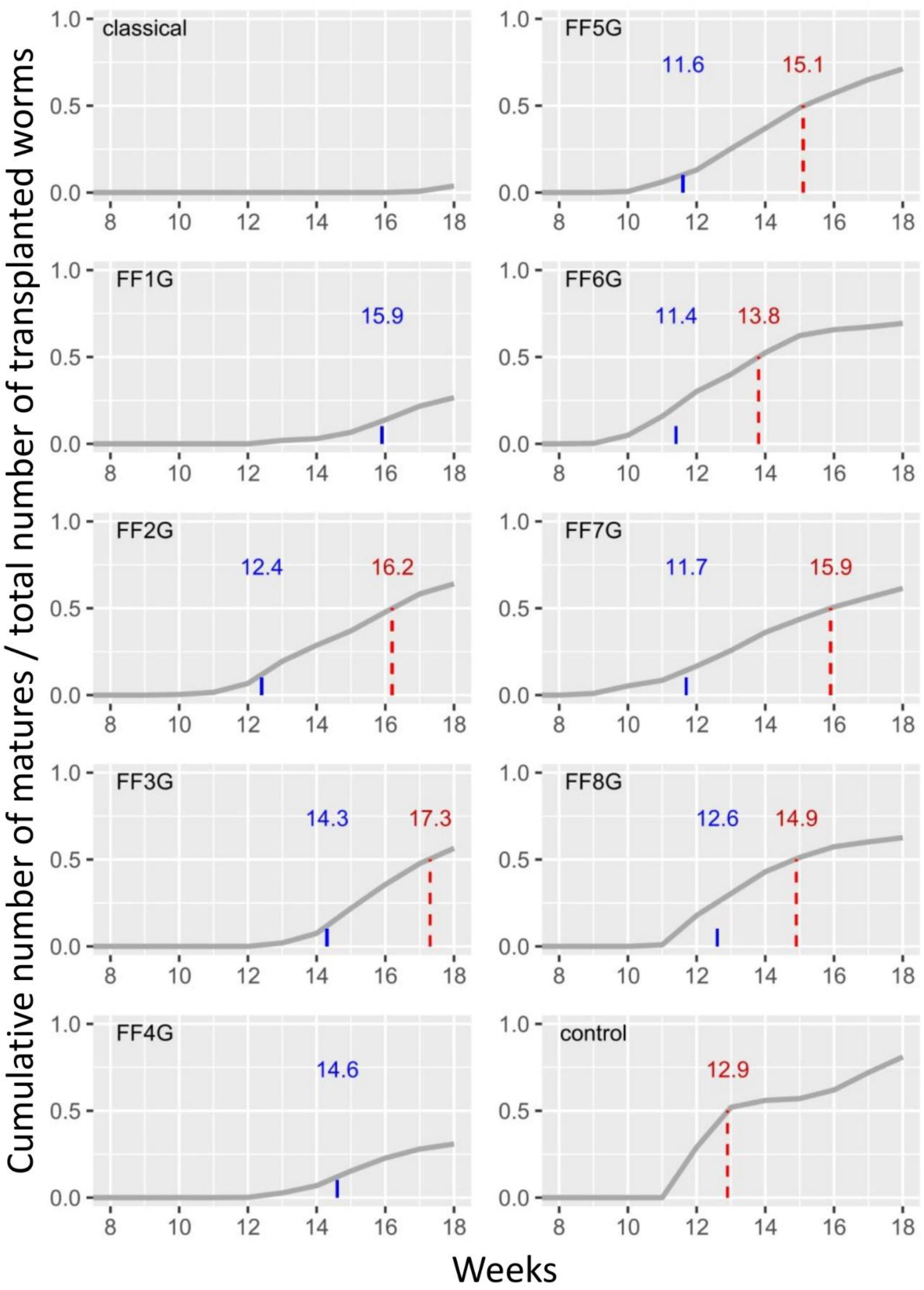
Cumulative plots of mature worms in each generation of the FF strain selection. “Classical” refers to the application of the typical food regime (Hauenschild & Fischer, 1969) to the original polymorphic strain. FF(1-8)G refers to the eight generations of selected worms . “Control” refers to the application of the final retained protocol to the polymorphic strain. The density, food regime and general statistics for each generation are described in Table 2. The dashed lines and numbers in red indicate when 50 % of the initially transplanted worms have matured (median maturation age). Numbers in blue indicate the average age of worms selected to spawn the entire next generation.

**Table 2.** Main statistics of the successive generations of the FF strain selection. Columns are named as in figure 4. The food regimes are (A) classical regime, 1^st^ month with 3ml microalgae twice per week, 2^nd^ month with 3ml Sera micron twice per week, 3^rd^ month with mixed Tetramin twice per week; (B), 1^st^ month with 3ml microalgae twice per week, 2^nd^ month with 1ml Sera micron twice per week, 3^rd^ month with mixed Tetramin/spinach as shown in Table 1; (C) Sera micron reduced to two weeks and spinach once per week from the 2^nd^ month, as described in Table 1.

|  | classical | FF1G | FF2G | FF3G | FF4G | FF5G | FF6G | FF7G | FF8G | control |
| --- | --- | --- | --- | --- | --- | --- | --- | --- | --- | --- |
| food regime | A | A | A | A | A | B | C | C | C | C |
| average density of boxes | 44 | 29 | 24.6 | 26.8 | 22.8 | 17.7 | 14.3 | 19.5 | 18.1 | 20 |
| number of boxes | 3 | 14 | 23 | 36 | 26 | 36 | 27 | 32 | 18 | 5 |
| total number of worms | 132 | 406 | 565 | 963 | 593 | 638 | 385 | 623 | 326 | 100 |
| total number of matures collected | 5 | 125 | 372 | 582 | 187 | 466 | 268 | 410 | 208 | 82 |
| average number of matures/box | 1.67 | 8.93 | 16.17 | 16.17 | 7.19 | 12.94 | 9.93 | 12.81 | 11.56 | 16.4 |
| number of matures collected/total % | 3.8 | 30.8 | 65.8 | 60.4 | 31.5 | 73 | 69.6 | 65.8 | 63.8 | 82 |
| (collected + dead)/total % | N/A | N/A | 97.9 | 69.4 | 79.3 | 93.3 | 97.7 | 94.2 | 97.9 | 95 |
| mortality % | N/A | N/A | 24.1 | 8.9 | 47.7 | 20.2 | 28.1 | 28.4 | 34 | 13 |
| sex ratio. % females | N/A | 41.6 | 51.7 | 46.7 | 55.1 | 52.1 | 46.4 | 48.8 | 52.9 | 51.2 |
| median mature age (weeks) | N/A | N/A | 16.2 | 17.3 | N/A | 15.1 | 13.8 | 15.9 | 14.9 | 12.9 |
| average age of next gen. parents (weeks) | N/A | 15.9 | 12.4 | 14.3 | 14.6 | 11.6 | 11.4 | 11.7 | 12.6 | N/A |

To determine whether genetic selection plays a role in fast cycling, we raised animals from the polymorphic stock using the same density and food conditions as those used in the last three generations of selected worms (FF6G-8G). The maturation curve of the polymorphic worms (Figure 2, control) does not appear significantly different from that of the selected worms FF6G-8G. The time of the first appearance of maturation for the polymorphic worms is 2 weeks later compared to FF6G-8G (Figure 4, Supplementary Figure 1), but the median age of maturation for the polymorphic worms is the youngest obtained in the study (figure 4). The maturation frequency of animals over time for the polymorphic and first generations of selected worms (FF1G-2G) displays two peaks at 4-week intervals, even without the application of an artificial moonlight regime (Supplementary Figure 1). Single waves of maturation are then observed in selected worms (FF3G-8G). The bimodal maturation in unselected worms could be linked to an intrinsic circalunar maturation rhythm (Zantke et al., 2013), but this is merely a hypothesis. The overall sex ratio in selected worms was not significantly different from parity (Table 2) but there is a significant tendency for earlier male maturation (Supplementary Figure 1), resulting in a slight excess of males in the first 3-4 weeks of mature collection. This discrepancy was not displayed in the control experiment, suggesting that it might be the result of genetic selection. This is again a hypothesis as the determinism of sex in *Platynereis* is not currently known. The earliest maturing individuals were 60 days old but did not produce viable offspring. The earliest mature animals involved in successful reproduction were 68-70 days old. Between 7 and 10 viable batches of eggs were selected for beginning the next generation of FF worms. The average age of the parents used for these maintenance reproductions decreased rapidly (Table 2 and Figure 4 in blue) and stabilized around 12 weeks (FF6G-8G).

**Table 2.**
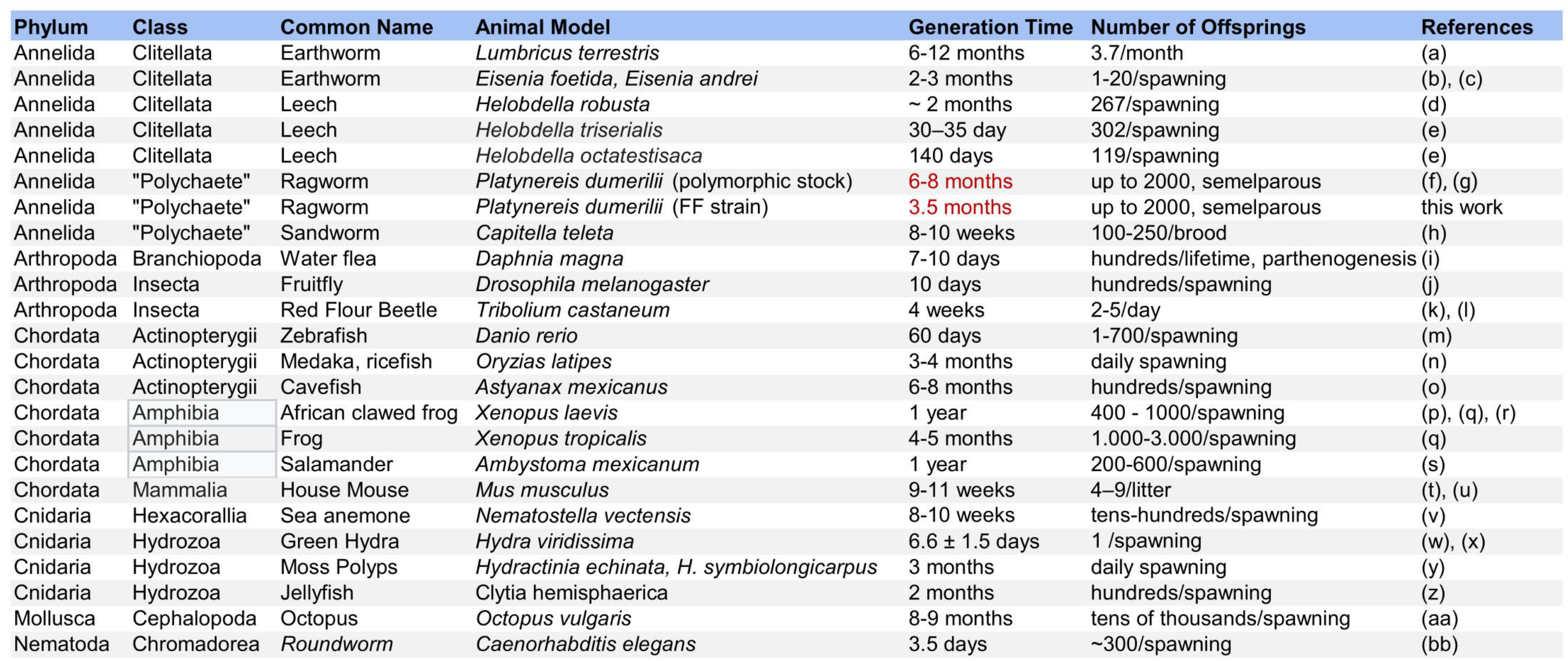
Reproductive characteristics of animal laboratory models used for transgenesis/genome editing experiments. References: (a) (Butt et al., 1992), (b) (Akazawa et al., 2021), (c) (Dominguez & Edwards, 2010), (d) (Shankland et al., 1992), (e) (Iyer et al., 2019), (f) (Williams & Jékely, 2016), (g) (A. H. L. Fischer et al., 2010), (h) (Seaver, 2016), (i) (Ebert, 2005), (j) (Yamaguchi & Yoshida, 2018), (k) (Schröder et al., 2008), (l) (Howe, 1962), (m) (Lawrence et al., 2012), (n) (Wittbrodt et al., 2002), (o) (Jeffery, 2020), (p) (Blackburn & Miller, 2019), (q) (Beck & Slack, 2001), (r) (Wolf & Hedrick, 1971), (s) (Tilley et al., 2022), (t) (Phifer-Rixey & Nachman, 2015), (u) (Weber & Olsson, 2008), (v) (Darling et al., 2005), (w) (Kaliszewicz, 2011), (x) (Massaro & Rocha, 2008), (y) (Frank et al., 2001), (z) (Lechable et al., 2020), (aa) (Iglesias et al., 2004), (bb) (Meneely et al., 2019)

We modified the density and food regime of the worm boxes over the two-year period, and unlike genetic selection, these two parameters appeared crucial in maintaining a fast-cycling culture, as explained below.

### Feeding and water change

The classical food regime utilized for our polymorphic *P. dumerilii* culture is based on the methods developed by Hauenschild and Fischer. Initially, we provided *Tetraselmis marina* microalgae obtained from a lab culture at the Institut Jacques Monod (courtesy of P. Kerner) to little worms aged from 4 days to 1 month old. This was followed by a suspension of Sera micron® for the second month, and then a mixed regime of Tetramin® and organic spinach from the third month onwards. These various food items are adapted to the mouth size of the fast-growing worms. To simplify the procedure, we first tried to eliminate the organic spinach. Table 2 provides a timeline of the successive regimes and quantities of food delivered to the selected worms.

During the selection process, we encountered two issues that significantly impacted the viability and maturation age of the worms, leading us to gradually modify our approach. The first issue was that some of the worms in certain boxes were unable to spin silk tubes. These worms would wander for the first two months and eventually assume a curled position with reduced mobility and feeding activity. This problem culminated in the FF4G generation, resulting in high mortality and a decrease in maturation (Figure 2). Changing the water did not correct this syndrome, and since it was not due to water fouling, we hypothesized that it might be caused by nutritional deficiency. As a result, we added ground spinach for the last four generations, and the abnormal behavior decreased from the 5th generation onwards. We then decided to provide spinach from the beginning of the second month along with Sera micron, as we reasoned that spinach must contain vitamins or other essential nutrients that are not present in Sera micron. This new approach eliminated the syndrome.

The second problem was mortality due to water fouling, which mostly resulted from feeding the worms with Sera micron. Initially, we provided quantities of the suspension that were too large for the small number of worms (15-30) in each box. However, Sera micron, which is primarily made with *Spirulina* cyanobacteria, is a highly nutritious food and very efficient for fast growth of the worms. Instead of eliminating it completely, we provided smaller quantities of the food suspension and limited the period of Sera micron to two weeks, as the worms quickly gained weight and could handle the larger particles of Tetramin. Initially, we attempted to raise the worms without performing time-consuming water changes to simplify the culture process. We believed that with a small number of worms, a controlled feeding regime, and the continued presence of dense populations of ciliates and flagellates in the boxes to clear unconsumed food, water fouling should be minimized. However, we reintroduced water changes at 60 days post-fertilization (dpf) to prevent water fouling in a few boxes that could have resulted in mass worm death. Water fouling typically results in yellowish and cloudy water. In contrast, the accumulated faeces do not seem to cause any problems for the worms despite the revolting look of old boxes. The box bottoms are also typically covered with green filamentous algae inoculated from the previous generation of boxes, which the worms occasionally feed on without issue (see below).

The last change we conducted related to food was replacing lab grown fresh *Tetraselmis marina* microalgae with commercially frozen microalgae of the same species (see Materials and Methods). We calculated a dilution factor for the microalgae that was equivalent to the dose of fresh algae we had been using. Half of the FF8G boxes were fed fresh algae, and the other half were fed frozen algae. We took care to compare boxes from the same parents. After 6 weeks of growth, we did not observe any significant differences in growth between the two types of food (Supplementary Figure 2). Therefore, the time-consuming task of culturing the microalgae can be replaced with relatively inexpensive commercial algae, given the small quantities used.

### Worm density

Worm density appears to be the most important factor for fast maturation. The traditional method for raising worms involves alternating between high-density and low-density boxes (Kuehn et al., 2019). Batches of small juvenile worms obtained from a single pair of worms in beakers are transplanted into one to three growth boxes, depending on the perceived density of worms. These worms will produce a dense layer of small tubes at the bottom of the box after a few weeks. However, their growth is slow, and maturation will not occur if the density remains above one hundred worms per box. Increasing food doses does not help because small worms seem to inhibit each other’s growth. Therefore, high-density boxes are transplanted into low-density boxes. The recommended density of worms for fast growth has typically been 30 worms per square box (around 500 individuals.m^-2^) (ref). Consequently, we decided to transplant low-density boxes directly from beakers containing 10 dpf feeding and growing little worms. Once food and water fouling problems were resolved, it became evident that the overall survival of these small, selected juveniles up to maturation age was excellent, approaching 100% in most cases, and their growth was very quick. To determine the optimal worm density for growth and maturation, we compared boxes of the FF6G generation that received different numbers of small worms (Table 2). Based on these data, we determined that the optimal density was 20 worms per box (300 individuals.m^-2^), significantly less than the previous conditions used.

## Discussion

As described above, the life cycle of *P. dumerilii* is lengthy and complex. This has hindered the widespread adoption of this model on a global scale. In response to this challenge, we propose in this article a new and fast-reproducing strain called “Fast-Forward” (FF). The “Three Rs” principle outlined in Directive 2010/63/EU of the European Union legislation reflects the scientific community’s current trend to Replace, Reduce, and Refine animal experimentation. The protocol presented in this article aims to reduce the resources required to culture *P. dumerilii* while accelerating and simplifying the management of its culture conditions. According to the Refinement principle of the “Three Rs”, we anticipate that the proposed modifications to the care practices will alleviate the distress experienced by the animal, thus reducing the variability of scientific results. Improving data quality indirectly

contributes to the Reduction principle, as fewer animals are required to obtain valuable results. However, the use of live animals remains a widely used strategy in developmental genetics, and it is an area that must be improved in the future.

### Suitability of the present culture method for different types of biological studies

The original laboratory culture of *P. dumerilii* was developed in 1953 in Germany by Carl Hauenshild and Albrecht Fischer (A. Fischer & Dorresteijn, 2004; Hauenschild & Fischer, 1969). Since then, all laboratories that have developed cultures appear to have used offshoots of this original German culture. This strain initially included a mixture of Atlantic and Mediterranean worms, which may partly explain the high level of genetic polymorphism. A complete culture method was posted online for a long time on the Platynereis.de website, which was cited in many previous articles and helped many groups establish their own culture. Although this website is no longer available, a copy of the method can be found on the Platynereis.com website. Several groups have since attempted to simplify the method. A recent effort has resulted in a “scalable culturing system” (Kuehn et al., 2019) for starting a small-scale culture without a dedicated thermostatic room. This method simplifies the food regime, alternating between *Spirulina* powder for the youngest worms and Sera micron for juveniles up to sexual maturation.

Our technique offers several steps for simplification, enabling the compaction of culture and preservation of wild-type stock and several different genetically modified strains in a limited space. We use flat boxes that can be stacked with at least two boxes on each shelf, without hindering gas exchange into the water. Piling up to four boxes has also been done without any issues. However, it is important to use plastic boxes with non-airtight lids (refer to Materials and Methods). We do not use air bubbling at any stage, due to the low-density of the culture and carefully managed food regime. Compared to earlier methods, our technique significantly reduces the need for water changes. With only one water change required when the worms reach two months (instead of every two weeks), a low-density box of worms can complete its cycle using only one litre of NFSW for a maximum of four months. This minimizes the workload, expenses, and natural resources required, making it possible to maintain bigger cultures in the future.

The main advantage of this culture technique is the acceleration of the reproduction cycle, which is a crucial step in establishing *P. dumerilii* as a valuable model for transgenesis and genome editing techniques. Measuring the average life cycle of *P. dumerilii* has seldom been done in the past, but previous methods have clearly resulted in much longer life spans and, consequently, reproduction cycles. The culture technique used at the Institut Jacques Monod for previous works since 2009, following the Hauenschild/Fischer method, involved using high-density boxes for an extended period (3-4 months) before worms were transferred to low-density boxes for maturation. As a result, very few worms matured before 4 months, and even after dispatching to low-density boxes, few worms matured before 5-6 months. The recently proposed scaled-down, cost-effective in-lab culture (Kuehn et al., 2019) does not focus on speeding up the cycle and maintains high-density boxes (more than 300 worms/box) for two months after fertilization. A work on *corazonin* effects on growth and maturation (Andreatta et al., 2020) mentions a median maturation time for control polymorphic worms of around 8 months, compared with 3.5 months with our FF strain and rearing protocol. In contrast to these lengthy periods of culturing at high density, the crucial parameters manipulated in the current technique are the immediate establishment of low-density boxes and the careful selection of fast-growing small worms at ten days.

This culturing technique may not be suitable for all experimental purposes. For instance, some teams study biological rhythms, especially those that are controlled by the lunar cycle (Poehn et al., 2022). The complete suppression of the artificial moon cycle and the maturation of most worms within a single moon cycle (28 days) are clearly inadequate for these types of studies. Others investigate time-related processes such as regeneration (Vervoort & Gazave, 2022) with recorded timelines under carefully controlled culture conditions. Using the FF strain and fast-cycling protocol, it will be necessary to test the capabilities of the selected worms in the specific process being studied and record new standard time courses accordingly. The elimination of high-density boxes means that the culture method does not allow for keeping large numbers of medium-sized juveniles, which may be impractical for teams working on processes occurring at these stages.

### Comparison with other metazoan models

Table 2 provides an overview of current metazoan models that have been acclimated in laboratory settings for experiments involving genetics and reverse genetics. This list includes a mix of models that have been developed for a long time, some for more than a century, with well-established protocols and numerous publications utilizing these techniques (such as the house mouse, fruit fly, and *Caenorhabditis elegans*), as well as several other models where efforts to develop genetic approaches are still in their early stages. Only a few models are available for studying the vast diversity of genetically dependent biological phenomena, as the duration of the life cycle is often a significant obstacle.

*D. melanogaster* and *C. elegans* are exceptions in this regard, as they depend on ephemeral food resources in the wild, specifically rotting fruits. As a result of selection pressure, they have evolved exceptionally fast development, growth, and sex maturation processes. However, these widely used models with fast life cycles are correlated with rapid evolution at the developmental, morphological, and molecular levels, making them “long evolutionary branch” models. Therefore, there is great interest in developing alternative model organisms with a short evolutionary branch for genetic investigations. These models can provide insights into the emergence of major anatomical features, cellular processes, and genetic machineries, simply because they are less derived in these respects compared to well established models.

The current list of short evolutionary branch metazoans includes notable species such as amphioxus (Louis et al., 2012), hemichordate worms (Simakov et al., 2015), *P. dumerilii* (Özpolat et al., 2021), the myriapod *Strigamia maritima* (Chipman et al., 2014), and the sea anemone *Nematostella vectensis* (Putnam et al., 2007), representing chordates, deuterostomes, annelids, arthropods, and cnidarians, respectively. Most of these species occupy key phylogenetic positions for reconstructing ancestral states in the metazoan tree. Phylogeny and evolution are not the only motivations for developing new genetic models, as some species exhibit unique derived characters that are worth exploring at the genomic/genetic level, such as the tardigrade *Hypsibius exemplaris* (Yoshida et al., 2017) and its ability to undergo anhydrobiosis.

Our new protocol places *P. dumerilii* in a better position for development as a high-performing genetic model in the future. The median life cycle obtained with this method (around 15 weeks) places *P. dumerilii* in a comparable position to the widely used fish model *Danio rerio* (zebrafish, table 2). Among annelids, the fast culturing of P*. dumerilii* places it on par in terms of culture conditions with other models, including both clitellates and “polychaetes” that have been developed in the past. While clitellates have interesting developmental features such as large size and easily micro-injectable teloblasts, they are derived annelids, adapted to freshwater or terrestrial lifestyles. Among “polychaetes”, *Capitella teleta* (Seaver, 2016) has emerged with many publications in the last two decades. It belongs to the large clade Sedentaria. Lineage tracing by micro-injection (Meyer et al., 2010) and genome editing by CRISPR-Cas9 (Neal et al., 2019) have been used successfully in this species. More recently, the other sedentarian *Owenia fusiformis* (Carrillo-Baltodano et al., 2021; Helm et al., 2016) has also been shown to possess valuable characteristics for diverse biological studies. *P. dumerilii* however displays two main advantages over these models: being an errantia of the family Nereididae, it is widely considered to possess a more ancestral anatomy than the sedentarians and the reproduction by epitokous swarming allows for complete control of the reproduction event and easy manipulation of thousands of eggs and embryos, obtained daily using the fast-cycling culture method described here.

### Culture size

The packing of boxes in incubators allows for easy scalability without the need to dedicate a thermostatic room of adequate size and shape for culture. The elimination of the lunar cycle and the rapid maturation of worms after 10 weeks are also important factors. There is no need for two rooms with alternating lunar cycles to obtain mature individuals over a monthly period. The maturation peaks will only depend on the age of the boxes. Mature worms will be used for three main purposes: colony maintenance, crosses with transgenic/CRISPR-edited worms, and micro-injection. We have generally been selecting mature worms aged less than 90 days to start the next generation of the FF population, and this selection is currently maintained. Worms older than 90 days are used for genetics and micro-injection. However, *P. dumerilii* still displays some unfavourable characteristics for genetic experiments, such as having 2n = 28 chromosomes, which limits the possibility of complex genetic combinations. Maintaining homozygous strains will be challenging as it requires two homozygous adults of opposite sexes on the same day, which in turn requires a large pool of worms for each strain. Therefore, the solution for keeping homozygous strains may lie in the development of sperm (or even embryo) freezing techniques (Olive and Wang, 1997). Transgenic lines expressing fluorescence as heterozygotes are maintained by crossing transgenic worms with wild-type FF individuals and selecting transgenic progeny under an epifluorescence microscope as early as 10 days post-fertilization. Our group in Paris has successfully maintained three different strains in this manner with only four boxes, each containing 20 worms, for each strain (Balavoine, unpublished).

In the current culture conditions, assuming an initial seeding of 20 worms per box and a conservative collection efficiency of 60%, we can calculate a productivity of approximately 1-1.2 worms/box/week over a 10-week period. There will be an unproductive period of 8-10 weeks following, as a new generation of boxes is growing. For teams that require wild-type worms throughout the year, which will be the case if multiple transgenic strains need to be maintained, the number of boxes will need to be doubled with a shift of 8-10 weeks between the seeding of low-density boxes. This shift can be easily achieved by maintaining batches of the same generation of small worms in high-density boxes (>300 worms) from 10 dpf on, fed exclusively with algae, for 8-10 weeks. The worms will remain very small until they are used to create new low-density boxes.

The current study may not represent the ultimate effort in establishing the most efficient culturing system for *P. dumerilii*, but it is a significant step towards reducing the variability in the age of sexual maturation. We believe that there is still much progress to be made in improving the food regime, particularly during the early settlement of the worms. One potential solution could be to draw inspiration from a recent small-scale culture published by Kuehn et al. (2019), where they reduced the food regime to two items, as opposed to the four items utilized in our present method. Another area for improvement could be the replacement of natural filtered sea water, which can be costly for facilities not located near the sea, with artificial sea water. Hopefully, our findings, combined with advancements in developing efficient transgenesis, will encourage more groups in the future to adopt this remarkable model animal for their own studies and establish their own culture of *P. dumerilii*.

## Acknowledgements

We thank the CNRS, the University Paris Cité and the Institut Jacques Monod for funding this research. In addition, the Balavoine group was financially supported by the Agence Nationale de la Recherche (grant PRCI TELOBLAST ANR-16-CE91-0007) and the Fondation ARC pour la recherche sur le cancer (grant LSP 190375). GG was supported by a Master student fellowship of the EUR G.E.N.E. graduate school (#ANR-17-EURE-0013) that is part of the Université Paris Cité IdEx #ANR-18-IDEX-0001 funded by the French Government through its “Investments for the Future” program. We want to thank P. Kerner for his help in microalgal supplies and tips. We acknowledge the staff of the animal facility of the institute J. Monod for help in worm husbandry.

## Contributions

ML, GG, and GB shared most of the worm husbandry, data collection, curation and analysis. RS and EC helped with worm husbandry. ML, GG and GB wrote the manuscript. GB designed the study. All figure hand drawings are made by GG.

---

**Supplementary figure 1.**
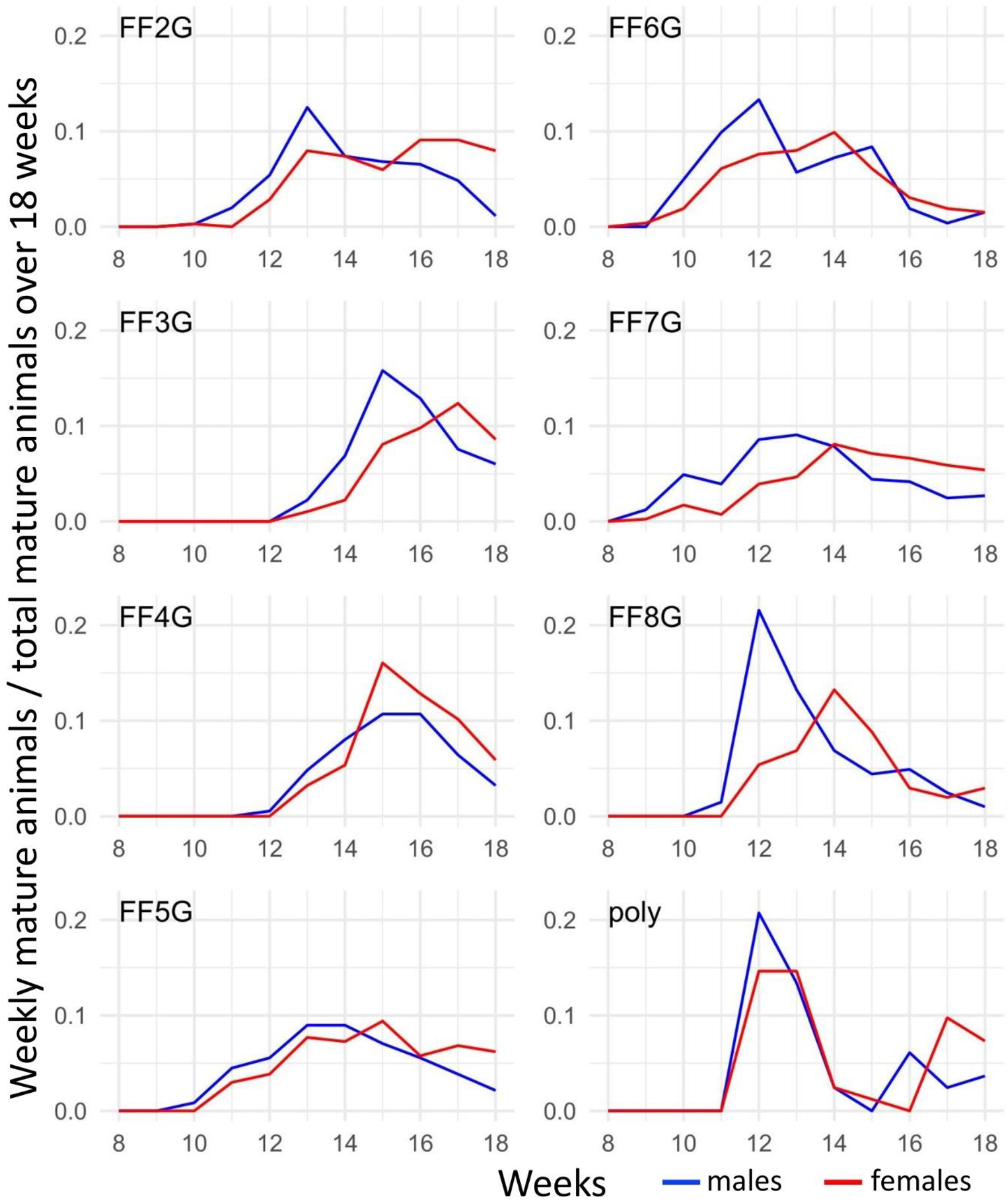
Sex ratio in mature animals across the selected FF generation. FF(2-8)G are the selected generation for which significant numbers of matures have been obtained. “poly” is the control culture with polymorphic worms raised with the same density and food regimen as the last three selected FF generations.

**Supplementary figure 2.**
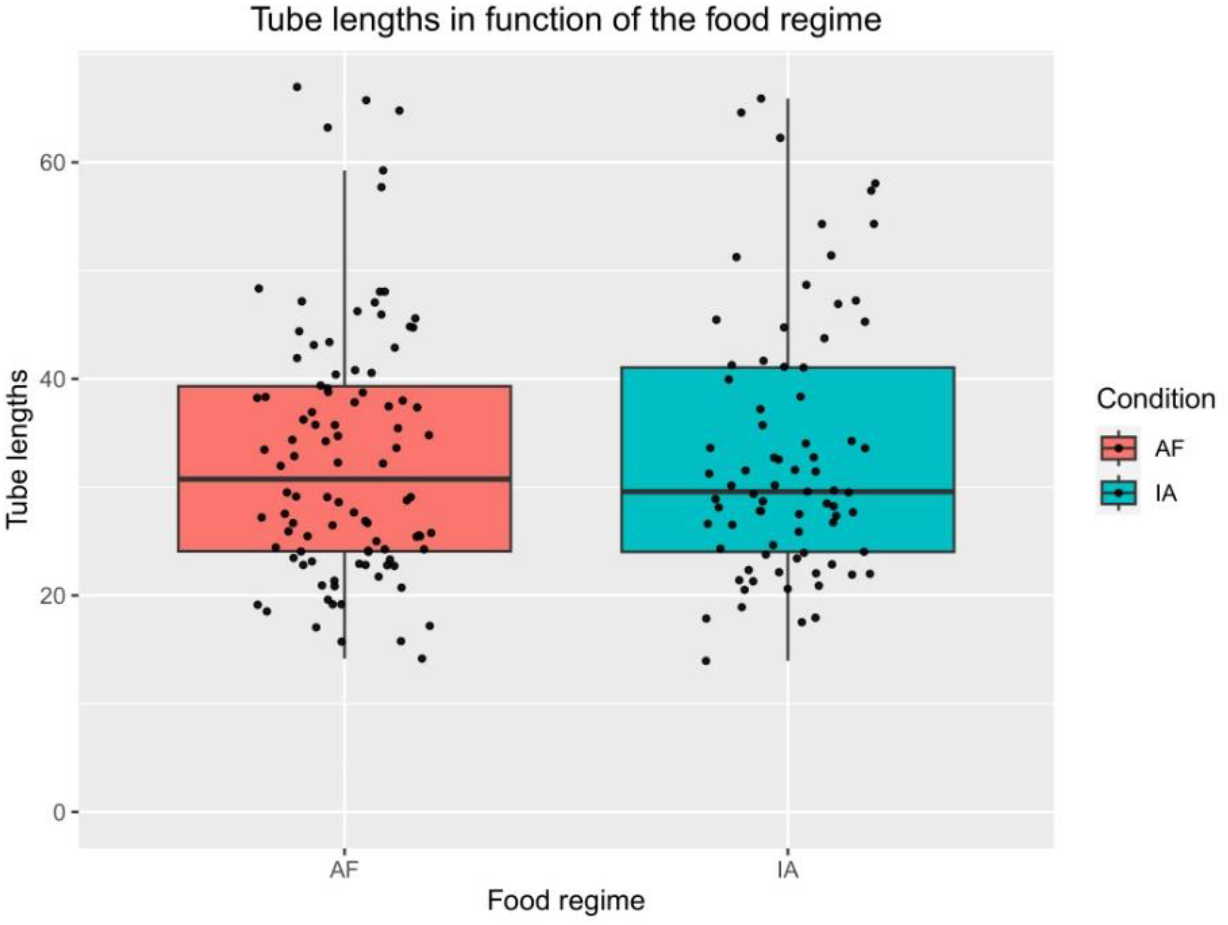
Growth in function of early algae food. Two alternative food regimes were applied on FF9G worms, one with one month of freshly grown microalgae (AF) and the other one with one month of frozen algae (IA, Instant Algae™) at two months post fertilization. As a proxy for growth, the length of silk tubes (closely representing worm length) covering the bottom of the boxes were measured using imageJ.

